# A Combined Immunophenotypic And Transcriptional Single-Cell Map Of First Trimester Human Fetal Liver Hematopoiesis

**DOI:** 10.1101/2021.12.29.474425

**Authors:** Mikael N.E. Sommarin, Rasmus Olofzon, Sara Palo, Parashar Dhapola, Göran Karlsson, Charlotta Böiers

## Abstract

Knowledge of human fetal blood development and how it differs from adult is highly relevant for our understanding of congenital blood and immune disorders as well as childhood leukemia, the latter known to originate in utero. Blood production during development occurs in waves that overlap in time and space adding to heterogeneity, which necessitates single cell approaches. Here, a combined single cell immunophenotypic and transcriptional map of first trimester primitive blood development is presented. Using CITE-seq (Cellular Indexing of Transcriptomes and Epitopes by Sequencing) the molecular profile of established immunophenotypic gated progenitors was analyzed in the fetal liver (FL). Classical markers for hematopoietic stem cells (HSCs) such as CD90 and CD49F were largely preserved, whereas CD135 (FLT3) and CD123 (IL3R) had a ubiquitous expression pattern capturing heterogenous populations. Direct molecular comparison with an adult bone marrow (BM) data set revealed that HSC-like cells were less frequent in FL, whereas cells with a lympho-myeloid signature were more abundant. Furthermore, an erythro-myeloid primed multipotent progenitor cluster was identified, potentially representing a transient, FL-specific progenitor. Based on the projection performed, up- and downregulated genes between fetal and adult cells were analyzed. In general, cell cycle pathways, including MYC targets were shown to be upregulated in fetal cells, whereas gene sets involved in inflammation and human leukocyte antigen (HLA) complex were downregulated. Importantly, a fetal core molecular signature was identified that could discriminate certain types of infant and childhood leukemia from adult counterparts.

Our detailed single cell map presented herein emphasizes molecular as well as immunophenotypic differences between fetal and adult primitive blood cells, of significance for future studies of pediatric leukemia and blood development in general.

## INTRODUCTION

Knowledge of our fetal blood system can facilitate generation of transplantable hematopoietic stem cells (HSCs) derived from pluripotent stem cells (PSCs) for future regenerative medicine (Wahlster and Daley, 2016), but also increase our understanding of congenital blood disorder and pediatric leukemia (Greaves, 2018).

Fetal and adult blood cells differ in cell cycle status, molecular profile as well as in composition of progenitor cells, and take place in different niches depending on the stage of embryonic development (Ivanovs et al., 2017; Kim et al., 2007; Roy et al., 2021; Yuan et al., 2012). Moreover, some innate immune cells are formed almost exclusively during fetal life (Ghosn et al., 2019). Much of our knowledge regarding embryonic and fetal hematopoiesis has been translated from studies of the murine system, but accumulating reports of the human counterpart complements the current model (Elsaid et al., 2020; Ivanovs et al., 2017). The blood system emerges in waves and is initiated by HSC independent hematopoiesis, first in the yolk sac and later also in the Aorta-Gonad-Mesonephros region (AGM) region, where immature erythroid and myeloid cells, critical for the growing embryo, emerge followed by erythro-myeloid and lympho-myeloid progenitors (Böiers et al., 2013; Ghosn et al., 2019; Ivanovs et al., 2017; Palis, 2016). At around day 27 (Carnegie stage (CS) 13) hematopoietic stem and progenitor cells (HSPCs) emerge in the AGM, and later migrate to the fetal liver (FL), where they mature and expand. The bone marrow (BM), the site of blood production in adults, starts to be colonized at the end of the first trimester, but the FL remains an active niche for hematopoiesis up until birth (Ivanovs et al., 2017).

Direct comparisons between human fetal and adult hematopoiesis have been challenging. The different waves and niches during development add to heterogeneity, and the sparsity of fetal samples makes studies in early development demanding, and differences are just beginning to be explored (Boiers et al., 2018; Notta et al., 2015; Roy et al., 2021). Recent studies from human FL hematopoiesis have shown that HSPCs become less proliferative with increased gestational stage, an observation that has been linked to HSPCs entering the fetal BM niche (Popescu et al., 2019; Ranzoni et al., 2020; Roy et al., 2021). Furthermore, HSPCs have been suggested to go through a change in lineage output during development, from oligopotency during fetal life to being dominated by mainly multi-or unipotent progenitors in adult (Notta et al., 2015) These temporal and spatial differences result in unknown heterogeneity calling for single cell analysis at different time-points of ontogeny.

Here, the transcriptome and immunophenotype of single first trimester FL cells were analyzed using CITE-seq (Cellular Indexing of Transcriptomes and Epitopes by Sequencing) (Stoeckius et al., 2017). In addition to demonstrating molecular changes of HSPCs related to developmental stage, immunophenotypic markers like CD123 (IL3R), CD135 (FLT3) and CD7 were observed to be more generally expressed, adding to heterogeneity when performing conventional direct comparisons based on immunophenotype. Using a nearest neighbor approach, FL HSPCs were directly compared to adult BM (Dhapola et al., 2021). This approach allowed for a comparison of progenitor composition within the fetal and adult HSPC compartment. In this analysis, molecular lympho-myeloid progenitors were found to be decreased from embryo to adult, whereas molecularly defined fetal HSCs were shown to increase with age. Importantly, by comparing common molecular signatures across cell types we could define a universal fetal core signature, with genes specifically up-or downregulated in fetal cells compared to adult. By using a publicly available RNA-seq data set from Acute Lymphoblastic Leukemia (ALL) patients (Gu et al., 2019), this fetal specific core signature was found to be enriched in some cases of pediatric leukemias, thus enabling separation of certain types of ALL based on the age of the patient.

Taken together, our combined single cell map highlights important molecular and immunophenotypic differences in human fetal and adult primitive blood cell development.

## RESULTS

### A transcriptional map of primitive cells from first trimester fetal liver

The emerging blood system is a complex mixture of cells from different niches and waves acting simultaneously, generating a heterogeneity that necessitates the use of single cell assays. In addition, immunophenotypic markers classically used to purify progenitors in cord blood (CB) and adult BM may not be equally expressed during development. To address these issues we analyzed FL HSPCs using CITE-seq, an unbiased, high-throughput single cell RNA-seq method, wherein information of immunophenotype is estimated simultaneously with the transcriptome, through oligo-barcoded tagged antibodies (antibody derived tags (ADTs))(Stoeckius et al., 2017). Early FL samples were obtained from aborted fetuses at first trimester, ranging in age from CS16 to 9 post-conceptional-weeks (pcw). HSPCs were selected based on expression of CD45 and CD34, and for lack of mature lineage markers (CD2, CD3, CD14, CD16, CD19, and CD235a) (*Figure 1a* and *Sup. Figure 1a*). The analysis focused on CS22, where 6732 cells from two embryos were captured for further analysis after quality control (*see method section*). Based on differentially expressed genes and molecular profile, 11 distinct cell states were identified (*Figure 1b-c*). The UMAP built showed a HSC cluster at the top, expressing genes like *MLLT3* (protein AF9) and *BST2*, the latter shown to be an activation-marker of HSCs in the murine system (Bujanover et al., 2018). Directly, subsequent to HSCs, cells with a Multipotent Progenitor (MPP)-like signature were located, followed by a distinct separation into a Megakaryocyte-Erythroid Progenitor axis (MEP, with genes like *GATA1* and *KLF1*), a Granulocyte-Monocyte Progenitor axis (GMP; with *MPO* and *CEBPD*) and a Lymphoid Progenitor axis (Ly-I/Ly-II). The lymphoid axis showed a gradual maturation from Ly-I to Ly-II with an increase in expression of *IL7R* and *LTB* among others. The Ly-II cluster was molecularly more primed towards the B lineage, expressing genes like *JCHAIN* and *VPREB1* (*Figure 1c*). The lymphoid clusters were located closest to a cluster with a DC-Mono (Dendritic Cell-Monocyte) progenitor signature and furthest away from the MEP axis. The Ly-III (Lymphoid Progenitor) was primed towards the T lineage with expression of *GATA3* and *CD3D* among others, but contained less than 60 cells, as did the DC-I (Dendritic cell precursor) cluster.

**Figure 1.**
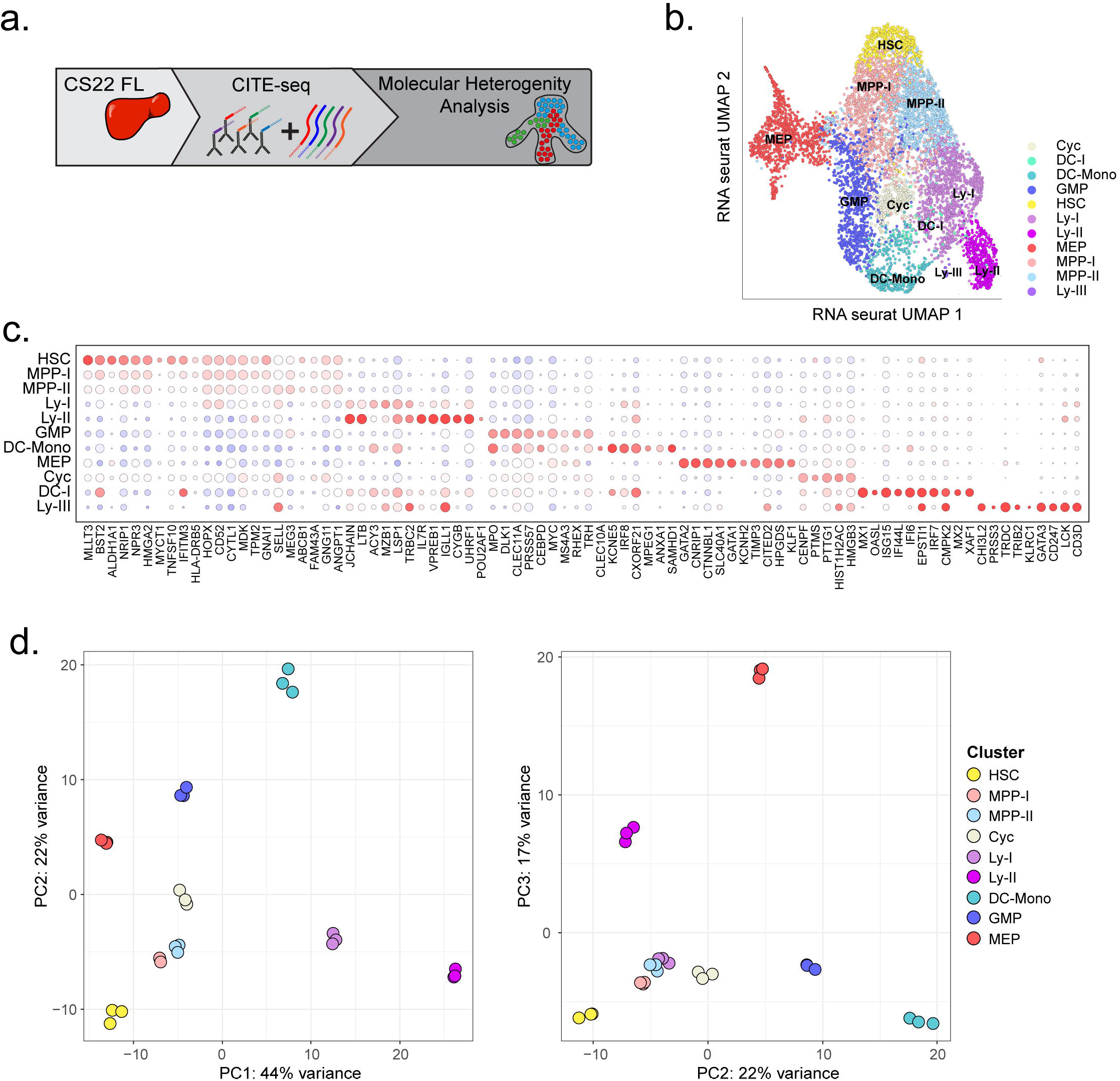
A transcriptional map of primitive cells from first trimester fetal liver. a) Schematic presentation of data in the figure. b) UMAP of cellular states within LIN^-^ CD45^+^CD34^+^ FL cells at CS22 (2 donors analyzed) c) Bead plot of differentially expressed genes between clusters, where the size of the circle is the fraction of cells with expression and the color represents the z-scored mean expression. d) Cells from the different cell states in b) were pseudo-bulked and PCA performed showing top 500 variably expressed genes. PC1 vs PC2 is shown to the left and PC2 vs PC3 to the right. Cluster name and color code label to the right.

To further investigate relatedness between the clusters, samples were pseudo-bulked and Principal Component Analysis (PCA) performed (*see method section*). The MPP-I and MPP-II were clustered together in the vicinity of HSCs as expected, whereas the Ly-II was positioned further away from the multipotent and myeloid clusters than Ly-I (*Figure 1d*). Thus, CITE-seq captured the molecular heterogeneity in early blood development and different progenitor populations could be defined based on expression of lineage-associated genes.

### Expression of immunophenotypic markers diverge through development

Next, the oligo-tagged antibodies (ADTs) were used to interrogate the conventional immunophenotypic populations within the HSPC compartment (*Figure 2a*). The HSC cluster expressed CD90 and CD49F, and lacked expression of CD38, all in agreement with markers used in conventional purifications of the HSC population (*Figure 2b*) (Majeti et al., 2007; Notta et al., 2011). Furthermore, CD201 (EPCR), a marker shown to define fetal HSCs, was also distinctly expressed in the molecular HSC cluster (Subramaniam et al., 2019). The comparatively immature lymphoid cluster (Ly-I) expressed CD10 and CD45RA, but was low in CD38 expression, indicative of Lympho-Myeloid Primed Progenitors (LMPPs) (Doulatov et al., 2010), whereas the relatively more mature lymphoid cluster (Ly-II) in addition to CD10 and CD45RA, expressed CD38, corresponding to a Common Lymphoid Progenitor (CLP) phenotype (Doulatov et al., 2010). This cluster also had distinct expression of interleukin 7 receptor (Il7R), known to mark CLP in the murine system (Kondo et al., 1997). The MEP cluster was positive for CD71 as expected (*Figure 2b*) (Mori et al., 2015; Notta et al., 2015). However, CD7, CD123 (IL3R) and CD135 (FLT3), the latter two markers traditionally used to define GMPs and Common Myeloid Progenitors (CMPs) (Doulatov et al., 2010; Manz et al., 2002), showed a broader expression pattern. The CD135 had a general surface abundance seen in all cell clusters, while CD123 was seen in myeloid as well as lymphoid clusters. CD7, on the other hand, was generally seen in the progenitor clusters, but low in the HSC cluster (*Figure 2b*).

**Figure 2.**
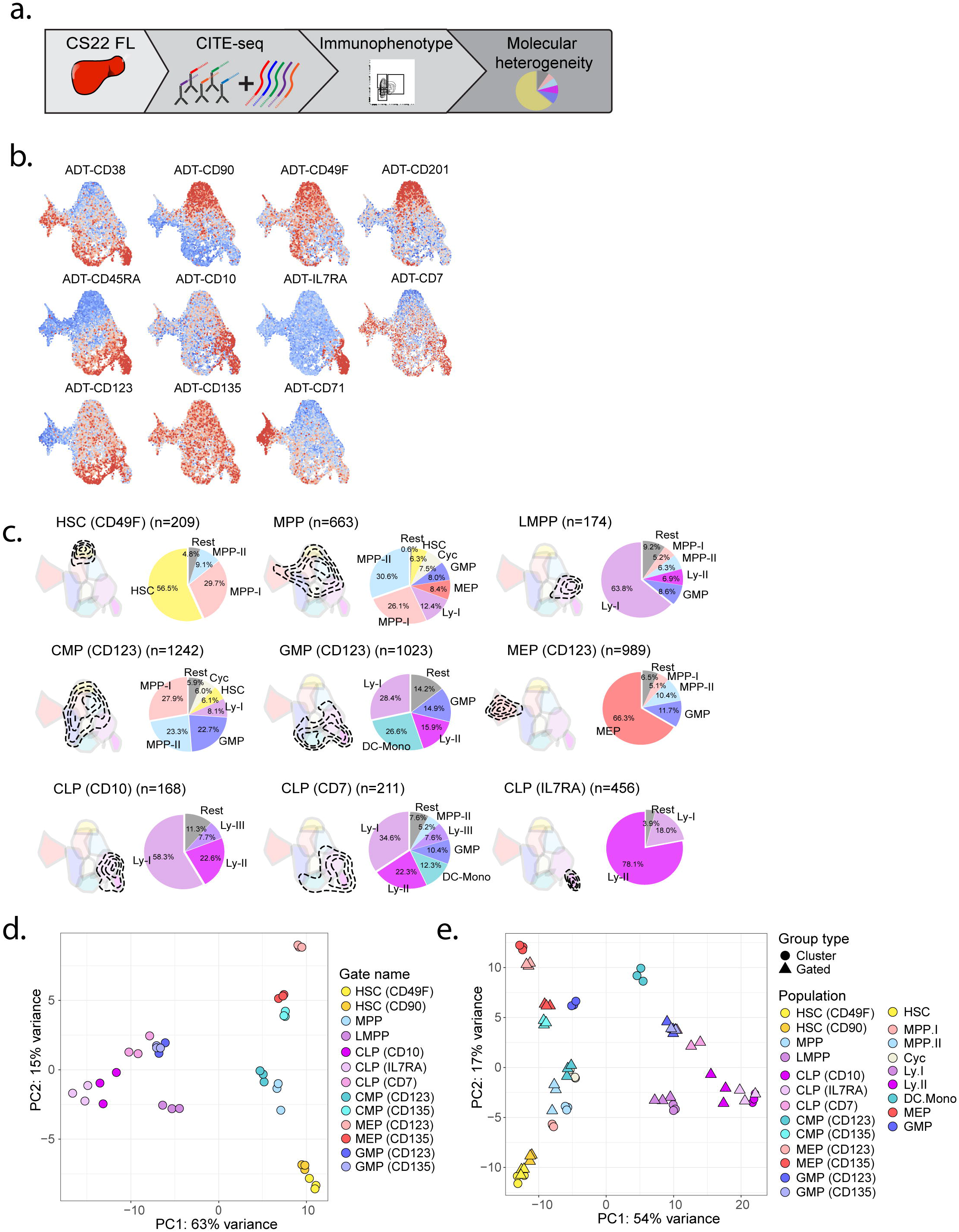
Expression of immunophenotypic markers diverge through development. a) Schematic presentation of data in the figure. b) FL CS22 UMAPs of ADT expression of single surface markers used for gating conventional cell populations in HSCPs, red represents high expression and blue low. c) Conventional immunophenotypic populations were gated using multiple ADT markers. Transcriptional cell states captured for each population are shown within the contours on the FL CS22 UMAP. Pie charts show percentage of the different molecular clusters within each immunophenotypic gated progenitor. Colors represent the cluster colors, defined in *Figure 1b*. d) PCA of ADT gated cell population of top 500 variably expressed genes. Cells were pseudo-bulked for the analysis (*see method section*). e) Combined PCA of ADT gated cell populations and the clusters defined in *Figure 1d*, showing top 500 variably expressed genes. (HSC^CD49F^: CD34^+^CD38^low^CD45RA^-^CD90^+^CD49F^+^; MPP: CD34^+^CD38^low^CD45RA^-^CD90^-^; LMPP: CD34^+^CD38^low^CD45RA^+^CD90^-^; CMP: CD34^+^CD38^+^CD10^−^CD45RA^-^CD123^+^, GMP: CD34^+^CD38^+^CD10^−^ CD45RA^+^CD123^+^; MEP: CD34^+^CD38^+^CD10^−^CD45RA^-^CD123^-^; CLP^CD10^: CD34^+^CD38^+^CD45RA^+^CD10^+^, CLP^CD7^: CD34^+^CD38^+^CD45RA^+^CD7^+^, CLP^IL7R^: CD34^+^CD38^+^CD45RA^+^IL7R^+^)

To define progenitor populations multiple surface markers are commonly used in combination. Thus, the ADT information was utilized to perform conventional immunophenotypic gating (*Sup. Figure 1b*). Molecular HSCs, MPPs, LMPPs and MEPs were well captured by the conventional adult immunophenotype, whereas CMPs and GMPs showed high heterogeneity as measured by the number of molecular clusters and their relative proportions captured within each immunophenotype (*Figure 2c* and *Sup. Figure 1c*). HSC gating (combining both CD90 and CD49F) resulted in over 50% purity of the molecular HSC cell state, and LMPPs defined as CD34^+^CD38^-^CD45RA^+^ yielded more than 60% of cells belonging to the Ly-I molecular state (Doulatov et al., 2010; Notta et al., 2011). On the other hand, immunophenotypic GMPs were heterogenous and 16% to 21% (using CD123 or CD135 gating respectively) of the cells molecularly belonged to the Ly-II lymphoid cell state (*Figure 2c* and *Sup. Figure 1c*)(Doulatov et al., 2010; Manz et al., 2002).

Conventional CLPs are typically defined based on expression of CD10, a marker also associated with priming towards the B cell lineage (Doulatov et al., 2010; Galy et al., 1995). Using this gating strategy, most of the captured cells were indeed part of a lymphoid transcriptional cluster, however almost 60% of the cells belonged to the lympho-myeloid Ly-I cell state, likely due to issues discriminating the cells based on CD38 expression. Therefore, we investigated whether other gating strategies could identify a transcriptionally more pure lymphoid cell state. In CB, CD7 has been shown to enrich for lymphoid progenitors, in combination with CD38 negativity (Hoebeke et al., 2007). However, CD7 surface expression measured with the ADT antibody was virtually absent in the CD38 negative fraction (*Figure 2b*). We therefore investigated if CD7 could capture a lymphoid cell state in the CD38 positive fraction. In the analysis, utilizing CD7 in combination with CD38 and CD45RA, 22% of the Ly-II cluster cells were captured, but more than 20% cells belonged to the GMP or DC-Mono cluster. Thus, this CD7^+^ ‘CLP-like’ population was heterogenous and mainly lympho-myeloid, in agreement with earlier study (Hoebeke et al., 2007). Next, IL7R, a marker that defines CLP in the murine system (Kondo et al., 1997), was utilized to investigate if it could define a more molecularly pure lymphoid cell state during fetal life. Importantly, when combining IL7R with CD45RA and CD38 almost 80% of the cells captured belonged to the Ly-II cell state, and the remaining cells were defined as Ly-I. Thus, IL7R was by far the best surface marker to capture the Ly-II cell state during early development (*Figure 2c)*.

To further investigate the heterogeneity between the immunophenotypic defined populations, a PCA plot of pseudo-bulked samples was made (*Figure 2d* and *Sup Figure 1d*). The CD10^+^ CLP population laid between the LMPP and the IL7R^+^ CLP, as our previous analysis indicated, whereas the ‘CLP’ gated on CD7 was located between GMP and CD10^+^ CLP. The myeloid CMP and MEP progenitors clustered differently depending on if CD123 or CD135 was utilized to define the population (*Figure 2c* and *Sup Figure 1c*). Next, the immunophenotypic populations were plotted together with the molecularly defined clusters from *Figure 1d*. Here, the immunophenotypic gated HSCs, MPPs, LMPPs and IL7R^+^CLP mapped to corresponding molecularly defined clusters, as well as MEP defined with CD123. The myeloid GMP and CMP populations were heterogeneous as expected from previous analysis, and the MEP and CMP molecular signatures differed depending on the gating strategy used (*Figure 2d-e* and *Sup Figure 1d-e*).

Thus, conventional immunophenotypic markers characterizing HSCs and LMPPs are largely conserved in the embryo, whereas immunophenotypic myeloid progenitors were molecularly heterogenous, but also markedly different depending on gating strategy used.

### Projection analysis defines a fetal-specific multipotent cluster with erythro-myeloid signature

Next, the fetal cells were directly compared with our adult BM data set of HSPCs, analyzed with CITE-seq in a similar way and using the same platform as the fetal cells (Sommarin et al., 2021) (*Figure 3a* and *Sup Figure 2a*). First, the pseudo-bulked immunophenotypic gated populations from fetal and adult were compared using PCA. PC1 separated the samples on developmental stage, which accounted for 53% of the variance (*Sup Figure 2b*). PC2 vs PC3 showed high heterogeneity between different progenitor types, developmental stage as well as the immunophenotypic markers used to define the progenitors, which makes immunophenotypic comparisons over ontogeny difficult. To preserve heterogeneity within the sample, but at the same time compare cells from different developmental stage and niche, a recently published nearest neighbor projection approach was used (Dhapola et al., 2021). A reference map was built, and the investigated cells mapped onto the reference, thus only molecular differences within the reference sample were considered. Cell composition can be determined and compared between samples based on the mapping and importantly, cell cycle effects were removed before the reference map was constructed. First using FL CS22 HSPCs as reference, 36% of the cells in adult BM mapped to the fetal HSC cluster, whereas only 7% belonged to this cluster in the FL CS22 sample. On the other hand, only 6% of the adult cells mapped to the lympho-myeloid Ly-I cluster, compared to 15% belonging to the Ly-I cluster in CS22 (*Figure 3b-c*). Thus, molecularly defined HSCs were enriched in adult BM compared to CS22 FL, whereas the Ly-I transcriptional cell state was reduced. Furthermore, MPP-I, which represents about 17% of the CS22 FL cells, were hardly detectable in the BM sample (2% of total cells). To narrow down the time window in development when these changes in cell states occurred, CB from the same data set as the adult BM, was analyzed, generating similar results as adult BM (*Figure 3b-c)* ((Sommarin et al., 2021). Next, two more embryonic time-points were analyzed from first trimester; CS16, and 9pcw (143 and 1139 cells analyzed respectively, each stage two donors). Here, 6 and 13% respectively of the fetal cells mapped to MPP-I, identifying MPP-I as a fetal specific cell state. These new analyses also confirmed that molecularly defined fetal HSCs increased with age from 1% at CS16 to 36% in adult BM, whereas Ly-I decreased from 27% in CS16 to about 6% in adult BM. At CS16, most cells mapped to lineage committed cell states in agreement with an active production of progenitors at this stage (*Figure 3b-d*).

**Figure 3.**
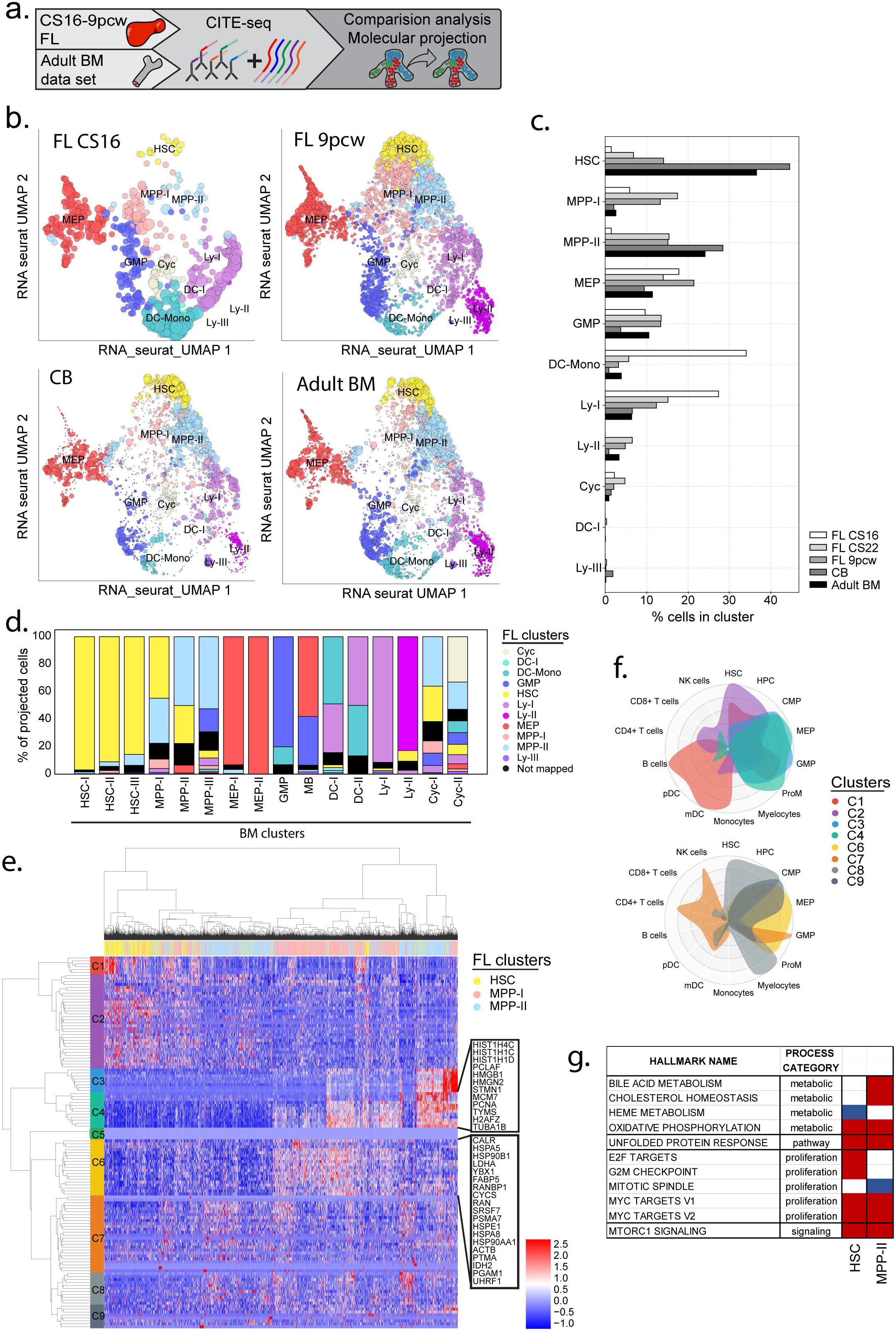
Projection analysis defines a fetal specific multipotent cluster with erythro-myeloid signature. a) Schematic presentation of data in the figure. b) Projection of FL CS16, FL 9pcw, CB and adult BM on the FL CS22 UMAP, size of dots represents mapping-score and color of dots represents FL derived clusters as in *Figure 1b*. c) Quantification of classified cells in all mapped developmental stages (FL CS16 143 cells, FL 9pcw 1139 cells, CB 6984 cells, adult BM 4905 cells). d) Classification of adult BM derived clusters on FL CS22. Colors represent FL CS22 derived clusters, and the x-axis represents BM derived clusters (*see also Sup Figure 2a*). e) Hierarchical clustering of primitive cell populations in FL CS22 (HSC, MPP-I and MPP-II) with genes defining the primitive populations. The genes were divided into 9 clusters (color coded). f) Cellradar plots of gene clusters in e (cluster names and color coded according to *Figure 3e*). g) GSEA using selected hallmark gene sets. MPP-I compared to HSC and MPP-II respectively. Color code according to NES (Normalized Enrichment Score) value; red; enriched in MPP-I, blue; downregulated in MPP-I. Selected gene sets with FDR q-value of <0.05 are shown.

Next adult BM was used as a reference, and FL cells from CS16 to 9 pcw were investigated. Intriguingly, almost no FL cells mapped to the adult HSC clusters and few to the more primitive MPP clusters, whereas the more mature MPP-III was enriched in the CS22 and 9pcw developmental stages. Furthermore, MEP-I was enriched in all stages of the FL, and at CS16 there was an increase of DC-I, corresponding to the DC-Mono cluster in the FL CS22 map (*Sup. Figure 2b-d*). The fetal specific MPP-I mainly mapped to MPP-III on the adult BM map (*Sup. Figure 2e*). Thus, transcriptionally defined adult HSCs were virtually absent in first trimester FLs.

To further narrow down the fetal specificity of the MPP-I cell state, we used a recently published data set that included samples from first and second trimester FL, as well as fetal, pediatric and adult BM (Roy et al., 2021). Mapping these data onto our CS22 UMAP showed enrichment of MPP-I in the early FL sample, while the fetal, pediatric and adult BM were almost completely depleted of the MPP-I cell state. Furthermore, there was a decrease in the proportion of MPP-I cell state from 8% to 3% from first to second trimester FL (*Sup Figure 2f*).

Next, differentially expressed genes among primitive clusters (HSC, MPP-I and MPP-II) were identified within CS22. In general, the MPP-I cell state appeared to be closely related to the HSC and MPP-II clusters, but with an upregulation of histone and heat shock related genes, amongst others (*Figure 3e, Sup. table 1*). Unfortunately, none of the ADT-antibodies were specific for the MPP-I, thus no surface marker in the panel could be used to purify the population (*data not shown*). The gene clusters identified were also investigated for lineage affiliation using CellRadar, a method where a gene set can be compared to public data of sorted hematopoietic populations (*see method section*). The clusters mainly associated with MPP-I (C4 and C6) showed an enrichment of an erythro-myeloid signature (*Figure 3e-f*). To further characterize MPP-I, Gene Set Enrichment Analysis (GSEA) was used, focusing on hallmark gene sets, where MPP-I differs from both HSCs and MPP-II (Liberzon et al., 2015; Subramanian et al., 2005). The significant gene sets of MPP-I were MYC targets, MTORC1 signaling as well as unfolded protein response and oxidative phosphorylation compared to both fetal HSCs and MPP-II, respectively, indicative of a more active metabolic state (False Discovery Rate (FDR) < 0.05 for all) (*Figure 3g*). Taken together, these data indicate that the MPP-I cluster harbors transient, FL specific erythro-myeloid primed multipotent progenitors.

### Cluster specific differential gene expression analysis defines a fetal-specific gene signature

The projection analysis performed identified clear differences in the composition of cell states and how it changes with developmental age. Furthermore, this analysis offers a unique possibility to directly compare gene expression differences between fetal and adult counterparts of the same cell types. Thus, to further understand how fetal and adult cell-types differ, the projected adult cells were compared to their fetal counterparts using DEseq2 (*Figure 4a*). Here, the differential gene expression analysis is performed on the whole set of genes, enabling identifications of differences in proliferation states. This is in contrast to the projection analysis, where cell cycle effects are removed to avoid interference in cell classification. Additionally, the clusters were pseudo-bulked, as with the immunophenotypic samples, and PCA performed (*Sup. Figure 3a-b*). Like the immunophenotypic samples, PC1 separated the clusters based on developmental stage. PC2 and PC3 separated samples depending on progenitor type and stage, where adult GMP, DC-Mono and Ly-II clustered further away from MPPs than fetal corresponding clusters, maybe indicative of these adult progenitors being more differentiated than their fetal counterparts (*Sup. Figure 3b*). In general, adult populations had more genes upregulated compared to fetal cells, and most differentially expressed genes were observed in HSCs (*Figure 4b* and *Sup. Figure 3c*). To investigate whether the transcriptional changes related to specific groups of genes, GSEA using hallmark gene sets was performed (Subramanian et al., 2005) focusing on gene sets involved in proliferation, apoptosis and immune system processes (Liberzon et al., 2015). Overall, pathways involved in proliferation were enriched in fetal progenitors, whereas gene sets involved in inflammation and immune system processes were downregulated, in agreement with an earlier study (Roy et al., 2021). The fetal specific MPP-I cells had only about 130 adult counterparts making the analysis less robust but showed upregulation of MYC targets in the fetal MPP-I, as was also detected when comparing it with the fetal HSC and MPP-II clusters (*Figure 3g* and *Figure 4c*).

**Figure 4.**
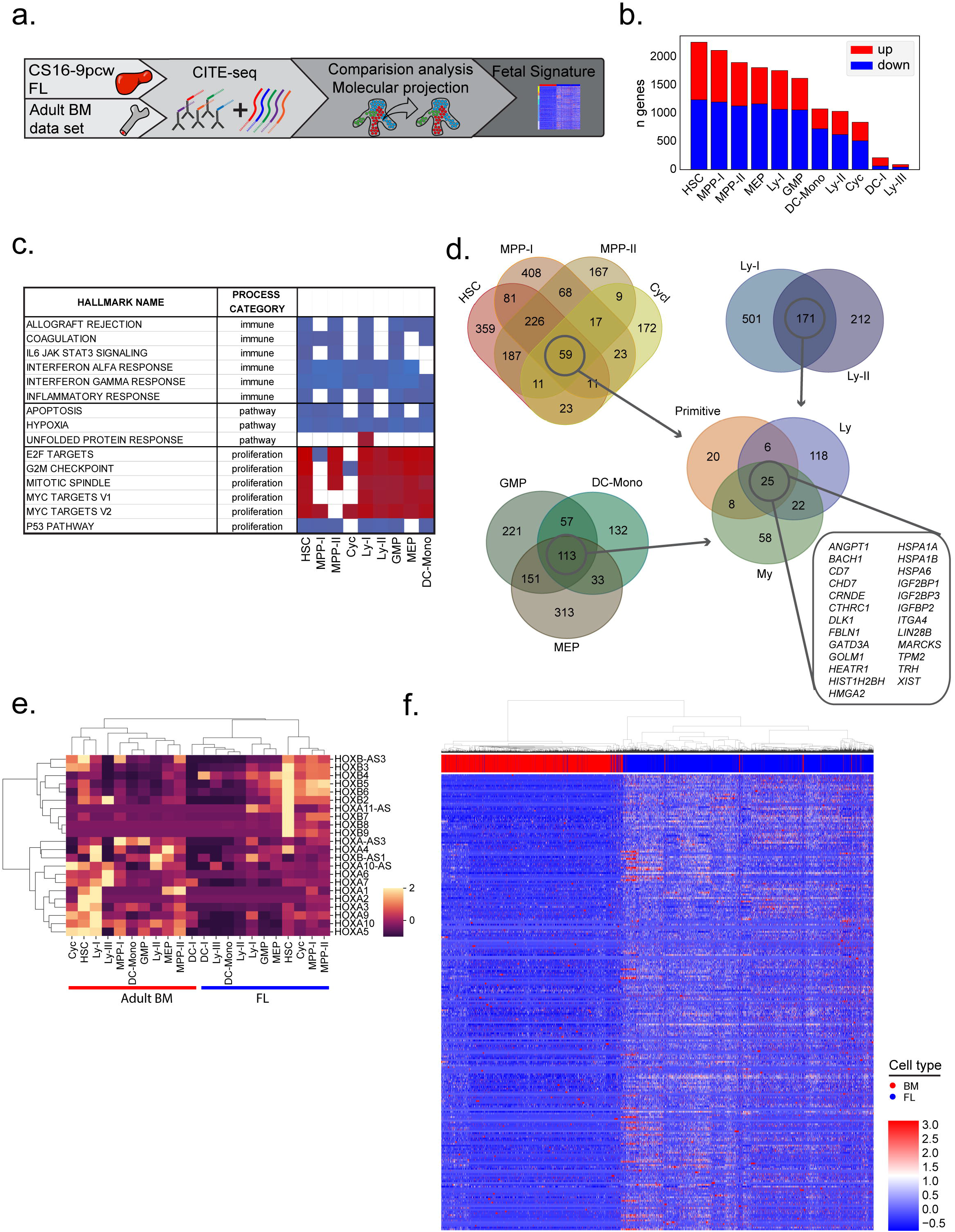
Cluster specific differential gene expression analysis defines a fetal specific gene signature. a) Schematic presentation of data in the figure. b) Number of up- and downregulated genes comparing adult BM cells mapping to the different FL clusters. Red bars; upregulated genes, blue bars; downregulated genes, adjusted p-value of <0.05. c) GSEA using hallmark gene sets involved in proliferation, selected pathways and immune processes, red; upregulated, blue; downregulated gene sets according to NES value. Selected gene sets with FDR q-value of <0.05 are shown. d) Venn diagrams of upregulated genes in FL compared to adult BM defined by fold change ≥2 and an adjusted p-value of <0.05 for each population. e) Heatmap of *HOX* gene expression per cluster in FL and adult BM. f) Heatmap of LIN^-^CD45^+^CD34^+^ adult BM (red) and FL (blue) cells, with cells displayed on the x-axis and fetal core genes on the y-axis.

Next, a Venn diagram was used to identify universal fetal specific up- and downregulated genes. A core set of 25 up- and 100 downregulated genes were identified (fold change ≥|2| and an adjusted p-value of <0.05, gender related genes were excluded in the subsequent analysis) (*Figure 4d* and *Sup. Figure 3d*). Among the top 25 upregulated genes *LIN28B, HMGA2, IGF2BP1* and *IGF2BP3* were found, all part of the LIN28B-let7 axis known to be involved in self renewal of fetal HSCs in the murine system (Copley et al., 2013) (*Figure 4d)*. Delta like gene 1 (*DLK1*) was also increased in all fetal populations, a gene shown to be a negative regulator of HSC formation in the mouse embryo (Mirshekar-Syahkal et al., 2013). *CHD7*, an epigenetic remodeler known to interact with RUNX1 and inhibit differentiation, was increased in all populations compared to adult (Hsu et al., 2020). Furthermore *CD7*, was more generally expressed among fetal progenitors, as was also seen at the protein level with the ADT marker (*Figure 2b*). Among downregulated genes human leukocyte antigen (HLA) complex of both class I and class II dominated, demonstrating a reduced antigen presenting capacity of the fetal immune system. *DNTT* (DNA nucleotidylexotransferase), known to induce diversity in the immunoglobulin chain in lymphoid progenitors, was also downregulated as expected in the fetal cells (Li et al., 1993). We also noticed that Homeobox (HOX) gene clusters were differently expressed in fetal and adult HSPCs, where *HOXB* genes were in general higher expressed in fetal and *HOXA* higher expressed in adult primitive clusters (*Figure 4e*).

Thus, by directly comparing fetal and adult counterparts from our projection analysis a fetal specific core signature was identified with genes specifically up- or downregulated in first trimester FL compared to adult counterparts (*Figure 4f*).

### The fetal gene signature differentiates between pediatric and adult ALL

By investigating neonatal blood spots and through twin studies, translocations giving rise to ALL in children could be backtracked to birth and an in-utero origin (Greaves, 2018). One question is if a remnant of the fetal signature can be detected in the pediatric leukemic cells. To investigate this, we first analyzed expression of the upregulated fetal core genes in a publicly available RNA-seq data set of an induced (I) PSC model of ETV6-RUNX1 (TEL-AML1) (Boiers et al., 2018). Differentiation of IPSCs to the B cell lineage recapitulates fetal lymphopoiesis and indeed many of the universal upregulated fetal genes were expressed in the IPS derived hematopoietic progenitors, as well as in the primary fetal progenitors investigated, but not in adult BM progenitors. The fetal genes were also to a large extent detected in the hematopoietic progenitors analyzed from the ETV6-RUNX1 expressing IPS cells (*Figure 5a*).

**Figure 5.**
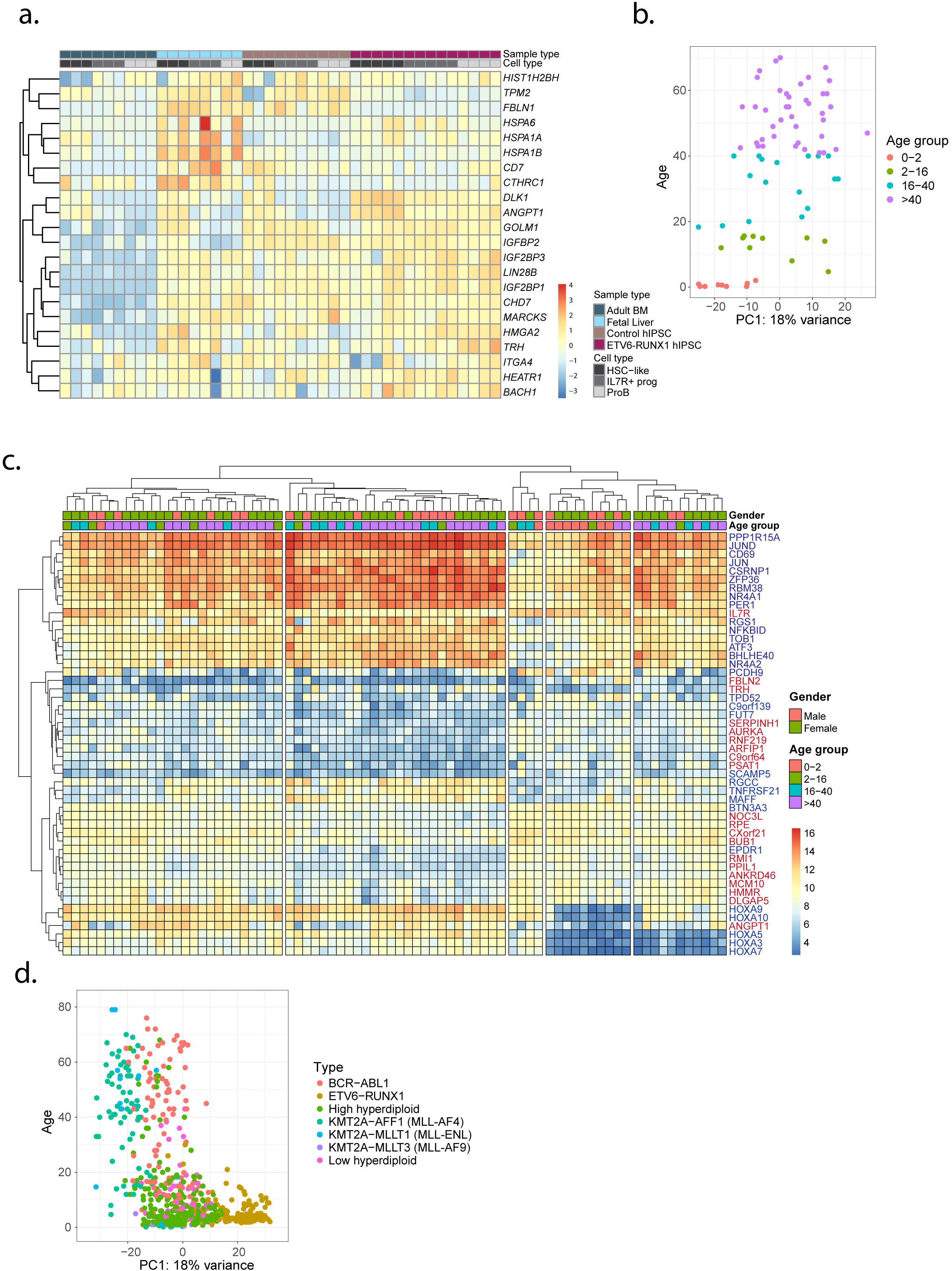
Fetal gene signature differentiates between pediatric and adult ALL. a) Heatmap based on FPKM values, displaying some of the universal fetal specific upregulated genes in a public available data set from an IPS model expressing ETV6-RUNX1. FL (CS17-22) and adult BM are shown as controls. Top rows describe the different sample types and cell types (‘HSC-like’, IL7R^+^ progenitor and proB). b) PCA of MLL-AF4 (KMT2A-AFF1) using the fetal gene signature, colors represent age groups. (red; 0-2 years, green; 2-16 years, turquoise; 16-40 years and purple; >40 years. PC1 vs age (years). c) Heatmap of the top 50 genes marking the highest and lowest values of PC1. Samples were hierarchically clustered into 3 groups, based on their gene expression. Top rows describe gender and age and to the right gene names are shown (Red label; upregulated in fetal cells, Blue label: upregulated in adult cells) d) PCA of B-ALL patient samples using the fetal gene signature, colour coded based on cytogenetic abnormality. PC1 vs age of patients. Colors describe translocation status with; *red*; BCR-ABL1, *orange*; ETV6-RUNX1, *dark green*; high hyperdiploid, *turquoise*; KMT2A-AFF1 (MLL-AF4), *blue*; KMT2A-MLLT1 (MLL-ENL), *purple*; KMT2A-MLLT3 (MLL-AF9) and *pink*; low hyperdiploid.

Next, using a publicly available RNA-seq data set with almost 2000 B-ALL patient samples, we aimed to investigate if the fetal core signature could be identified in B-ALLs of known in utero origin (Gu et al., 2019). As different RNA-seq methods were used in the study, B-ALL with different driver mutations of interest were selected, pooled and batch corrected (*Sup Figure 4a*). First, Mixed Lineage Leukemia gene (MLL1/KMT2A) fused with AF4 (AFF1) t(4;11) was investigated. MLL-AF4 is an initiating mutation that almost exclusively gives rise to B-ALL and is found in many different age groups, including infants, for whom a clear in utero origin has been demonstrated (Gale et al., 1997). A PCA plot was generated using the fetal core signature (all fetal up- and downregulated genes identified in primitive, lymphoid and myeloid progenitors; a total of 257 up and 526 down, gender associated genes were removed in the analysis) (*Figure 4d* and *Sup Figure 3d*). Intriguingly, the PC1 showed distinct separation based on age, discriminating between pediatric and adult samples (*Figure 5b* and *Sup Figure 4b*). A heatmap of the top 50 differentially expressed genes on PC1 identified a difference in expression of *HOXA9* and *HOXA10* between infants and adult, where most infants had lower expression of these *HOX* genes. *HOXA3, HOXA5* and *HOXA7* were also expressed at low levels in infants, as well as in some adult MLL-AF4 fusion ALL (*Figure 5c* and *Sup Figure 4c*). Indeed, a correlation has been observed between *HOXA* expression, prognosis and age in an earlier study, investigating *HOX* gene expression in ALL compared to normal progenitors (Starkova et al., 2010).

Next, more subtypes of ALL were investigated using the same fetal core signature. The PC1 again correlated with age, where a clear association was seen with ETV6-RUNX1 and hyperdiploid ALL, the two most frequent cytogenetic abnormality in children (Greaves, 2018), though for some of these leukemia types there were almost no adult patients represented in the material. In contrast, the adult associated translocation BCR-ABL, showed almost no spread along PC1 (Bernt and Hunger, 2014)(*Figure 5d* and *Sup Figure 4d-e*). Thus, in some subtypes of leukemia associated with children and in utero origin, part of a fetal core signature remains in the leukemic cells. The data have the potential to explain differences in pediatric and adult leukemia carrying similar driver mutation, potentially important for prognosis and therapy response.

## DISCUSSION

The hematopoietic system is a paradigm hierarchical organization of tissues. The structure has been built on careful immunophenotypic isolation and characterization of progenitors, which has formed the foundation of the hematopoietic hierarchy of today in both mouse and human (Jacobsen and Nerlov, 2019; Seita and Weissman, 2010). Recent developments in single cell omics have allowed for further purification and analysis of molecular heterogeneity, and with these findings the hematopoietic hierarchy can be regarded as a continuum rather than a step wise hierarchical organization (Laurenti and Göttgens, 2018; Velten et al., 2017). Until recently these studies have mainly investigated adult hematopoiesis, but more reports looking into human blood development are rapidly emerging (Jardine et al., 2021; Popescu et al., 2019; Ranzoni et al., 2020; Roy et al., 2021). So far these studies mainly focused on molecular heterogeneity, and it has until now been unclear to what degree the immunophenotype of CB and adult BM progenitors translate to early FL. Within this study we interrogated FL HSPCs using single cell RNA-seq, relying on CITE-seq to also capture immunophenotype together with the transcriptome. By focusing on early primitive hematopoiesis and by using CITE-seq the whole HSPC population could be interrogated and the relative proportions of all CD34 positive populations compared across developmental time-points, hereby also capturing the immunophenotype, enabling a unique view of the classically defined progenitors.

While most of the conventional markers of HSPCs (CD90, CD45RA, CD71 and IL7R) showed good correlation to the transcriptionally defined clusters, markers of GMPs and CMPs i.e., CD123 and CD135 (FLT3) were more generally expressed. This was even more apparent when utilizing the ADTs to perform immunophenotypic gating, which showed substantial molecular heterogeneity within GMPs and CMPs. The discrepancy in expression of surface markers during development thus may add to molecular heterogeneity in comparisons based on immunophenotype. Our data also shows that gating with CD123 or CD135 respectively for myeloid progenitors (CMP and MEP) captures molecularly heterogenous cell types (*Figure 2c* and *Sup Figure 1c*) (Doulatov et al., 2010; Manz et al., 2002). This also applies for the lymphoid surface markers, where CD10, mainly captured the lympho-myeloid cluster Ly-I, whereas our data revealed that IL7R in combination with CD38 and CD45RA could capture the Ly-II cell state with high purity in the embryo. This was opposite as to what was seen in adult where the population captured by CD10 comprised 70% Ly-II cells vs 43% when using IL7R (*data not shown* and (Sommarin et al., 2021)). Thus, CITE-seq detects molecular heterogeneity within immunophenotypic progenitors, but also identifies surface markers that are preserved throughout development.

An advantage with high-throughput single cell RNA-seq methods is that there is no need to pre-select populations based on immunophenotype. Instead, a broad selection of cells can be investigated, like in our case HSPCs. However, difficulties remain when cells from different stages and niches are compared. Earlier studies have merged different stages onto the same analysis, relying on batch correction to enable sample integration (Jardine et al., 2021; Popescu et al., 2019; Roy et al., 2021). However, when merging cell populations from different stages onto the same analysis, the developmental differences may take over and heterogeneity within the sample itself may be lost. The projection approach used herein allows for signals responsible for lineage determination to be kept, and additionally, by removing cell cycle effects through regression, avoiding differences in cell cycle status from disrupting the cluster definitions (Dhapola et al., 2021). By mapping the test cells onto the reference map, only molecular differences that separate the reference cells will be investigated. From this analysis, cell state differences over development could also be captured. As expected, major changes in cell states during development could be observed, with HSCs being increased with age, while the relative fraction of lymphoid progenitors (Ly-I and Ly-II) were decreased, in agreement with an earlier study (Roy et al., 2021). Furthermore MPP-I was found to be almost exclusively a fetal cell state, suggesting that this population could constitute a fetal specific progenitor population, arising from a HSC independent wave. The MPP-I cells displayed an erythro-myeloid gene signature, while maintaining a primitive gene program. Future studies are likely to investigate whether this population represent the erythro-myeloid progenitor described in mouse developmental hematopoiesis, originating in the yolk sac prior to definitive HSC formation (Ghosn et al., 2019; Palis, 2016).

Further analysis of the molecular differences between the FL and adult BM using differential gene expression analysis showed substantial ontogeny-dependent transcriptional differences. By utilizing the projection analysis to define molecularly similar cells, comparisons without interfering signals caused by differences in heterogeneity could be performed. This analysis showed that the HSCs experienced most transcriptional differences related to age. GSEA analysis grouped these differences into increased proliferation and reduction in inflammation compared to adult, in agreement with a recent study (Roy et al., 2021). Also, expression of HLA complex of both class I and class II are reduced in the fetus. HLA-B was found reduced in HSCs/MPPs in an earlier study, but we demonstrate a reduction of several different HLA classes in fetal HSPCs, all indicative of a reduced antigen presenting ability of fetal cells in general (Popescu et al., 2019).

The projection analysis of FL CS22 onto adult BM reference also revealed substantial molecular differences between molecular HSCs in FL and adult BM. The molecular FL HSCs mainly projected to adult BM MPP-I and MPP-III, but not to the adult HSC molecular clusters, even though cycling factors had been removed in this analysis. Thus, our data shows that multiple factors differ fetal and adult HSCs from each other, of general interest to be able to generate transplantable HSCs from differentiated PSCs in the future (Wahlster and Daley, 2016).

Childhood B-ALL has in many cases been shown to have a fetal origin (Greaves, 2018), and the leukemic cells could potentially retain expression of fetal specific genes or lack expression of adult specific genes (Symeonidou et al., 2021). By comparing the up- and downregulated genes in fetal and adult cells in each cluster, a core set of genes linked to fetal or adult identity were identified. These genes, together with genes linked to fetal and adult primitive, lymphoid and myeloid identity, were used to investigate childhood leukemia from publicly available data of almost 2000 B-ALL samples with different cytogenetic abnormalities (Gu et al., 2019). The fetal core signature correlated in same cases with age of the patient, and for MLL-AF4 it was clear that *HOXA* genes were downregulated in many infants compared to adult, in agreement with an earlier study, where a correlation was observed between lower expression of *HOXA* genes, poor prognosis and young age (Starkova et al., 2010). Of note in our fetal-adult data set *HOXA* genes were specifically lower expressed in fetal HSCs compared to adult (*Figure 4e*).

Our study links immunophenotype with transcriptome at the single cell level, providing a unique map of human fetal blood development. Comparison to adult hematopoiesis gives insight into how the hematopoietic progenitor compartment changes with development. A fetal core signature with universal up-or downregulated genes depending on developmental stage could be identified, a gene set that could be used to separate certain types of B-ALL samples based on age. The importance of the fetal core genes in the formation of the pre-leukemic clone in utero and in leukemia progression and development in general remains to be explored.

## SUPPLEMENTAL FIGURES

**Sup. Figure 1.**
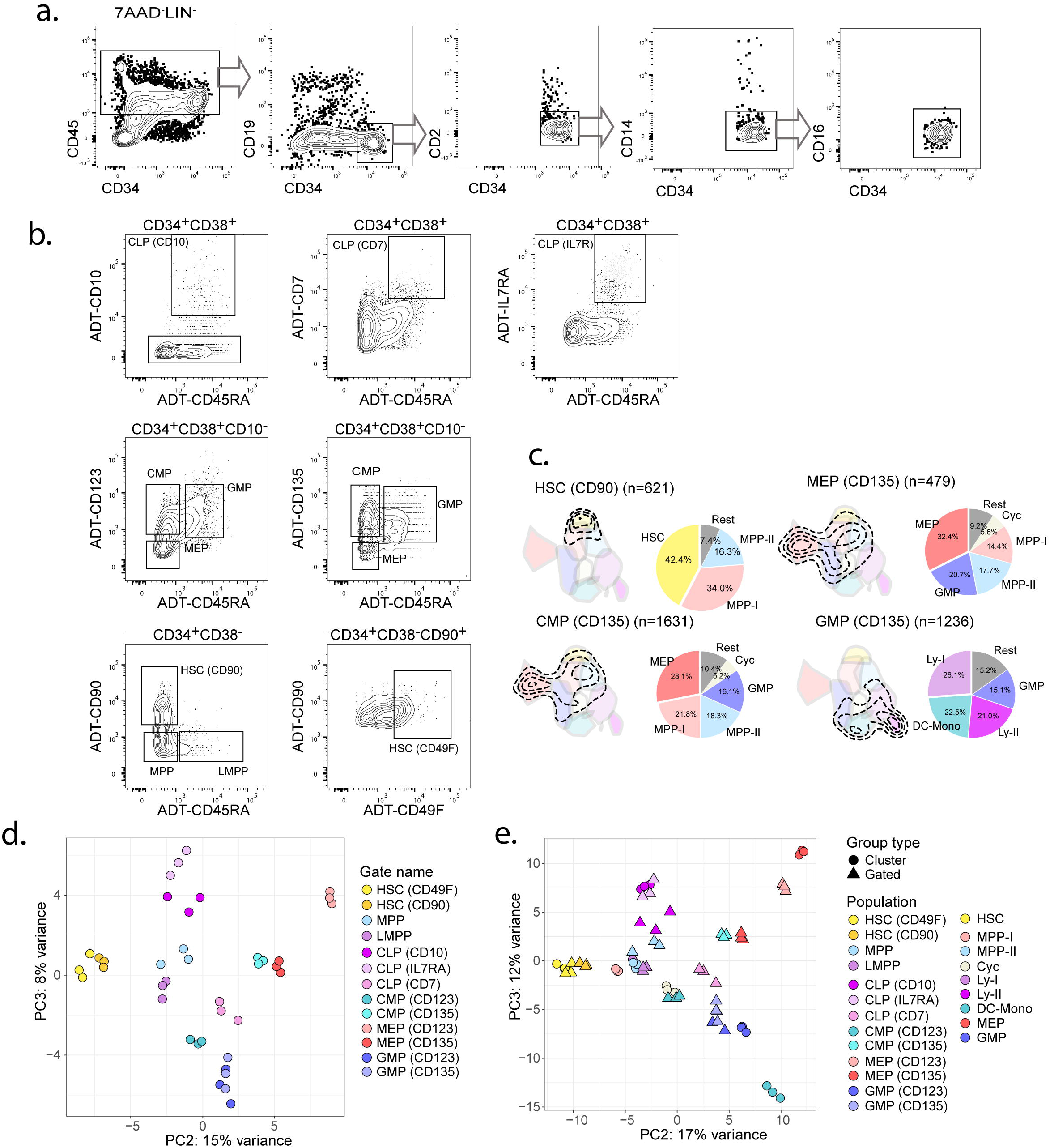
Purification of primitive FL cells and immunophenotypic gating using ADTs. a) Sorting strategy of CS22 LIN^-^CD45^+^CD34^+^ FL cells. Viable cells (7-AAD^-^) were selected based on size (scatter) and gated negative for mature lineage markers (CD3, CD235a). Further gating is indicated in the figure. b) Gating of conventional immunophenotypic populations using ADT signals. Top row CLP gating, middle row myeloid progenitors (CMP, GMP, MEP) and bottom row primitive populations (HSC, MPP, LMPP). c) Conventional immunophenotypic populations were gated using multiple ADT markers. Cell states captured for each population are shown within the contours on the FL CS22 UMAP. Pie charts show percentage of the different molecular clusters within each immunophenotypic gated progenitor (colors represent the cluster colors, as in *Figure 1b)*. d) PC2 vs PC3 of ADT gated cell populations of top 500 variably expressed genes. Cells were pseudo-bulked for the analysis. e) Combined PCA of ADT gated cell populations and the cell states defined in *Figure 1d*, of top 500 variably expressed genes. PC2 vs PC3 are shown. (HSC^CD90^: CD34^+^CD38^low^CD45RA^-^CD90^+^; CMP^CD135^: CD34^+^CD38^+^CD10^−^CD45RA^-^CD135^+^, GMP^CD135^ CD34^+^CD38^+^CD10^−^CD45RA^+^CD135^+^; MEP^CD135^: CD34^+^CD38^+^CD10^−^CD45RA^-^CD135^-^)

**Sup. Figure 2.**
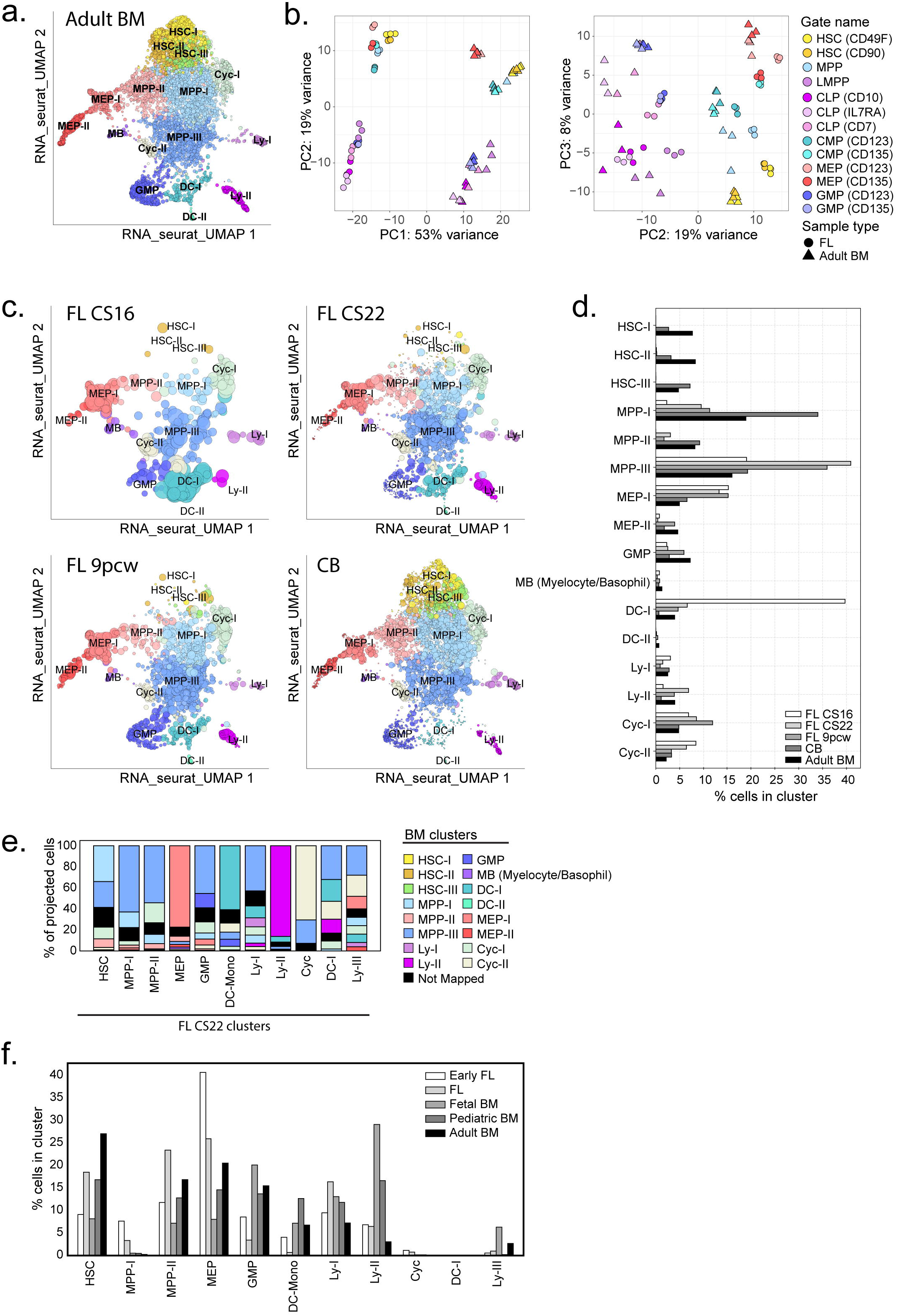
Projection of primitive FL and CB cells on adult BM reference. a) UMAP of LIN^-^CD45^+^CD34^+^ adult BM cells with cluster annotations from (Sommarin et al., 2021) b) PCA of pseudo-bulked immunophenotypic populations from FL CS22 and adult BM of 500 top variably expressed genes. PC1 vs PC2 left and PC2 vs PC3 right plot. c) Projection of FL CS16, FL CS22, FL 9pcw and CB on the adult BM UMAP. Size of dots represents mapping-score and color of dots represents BM clusters. d) Quantification of classified cells in all mapped developmental stages. e) Classification of FL CS22 derived clusters on adult BM, colors represent BM derived clusters according to *Sup. Figure 2a*, and the x-axis represents FL CS22 derived clusters. f) Quantification of classified cells in all mapped developmental stages from (Roy et al., 2021). In total 90-95% of the cell from the different stages mapped. (Early FL; first trimester, FL and fetal BM; second trimester.)

**Sup. Figure 3.**
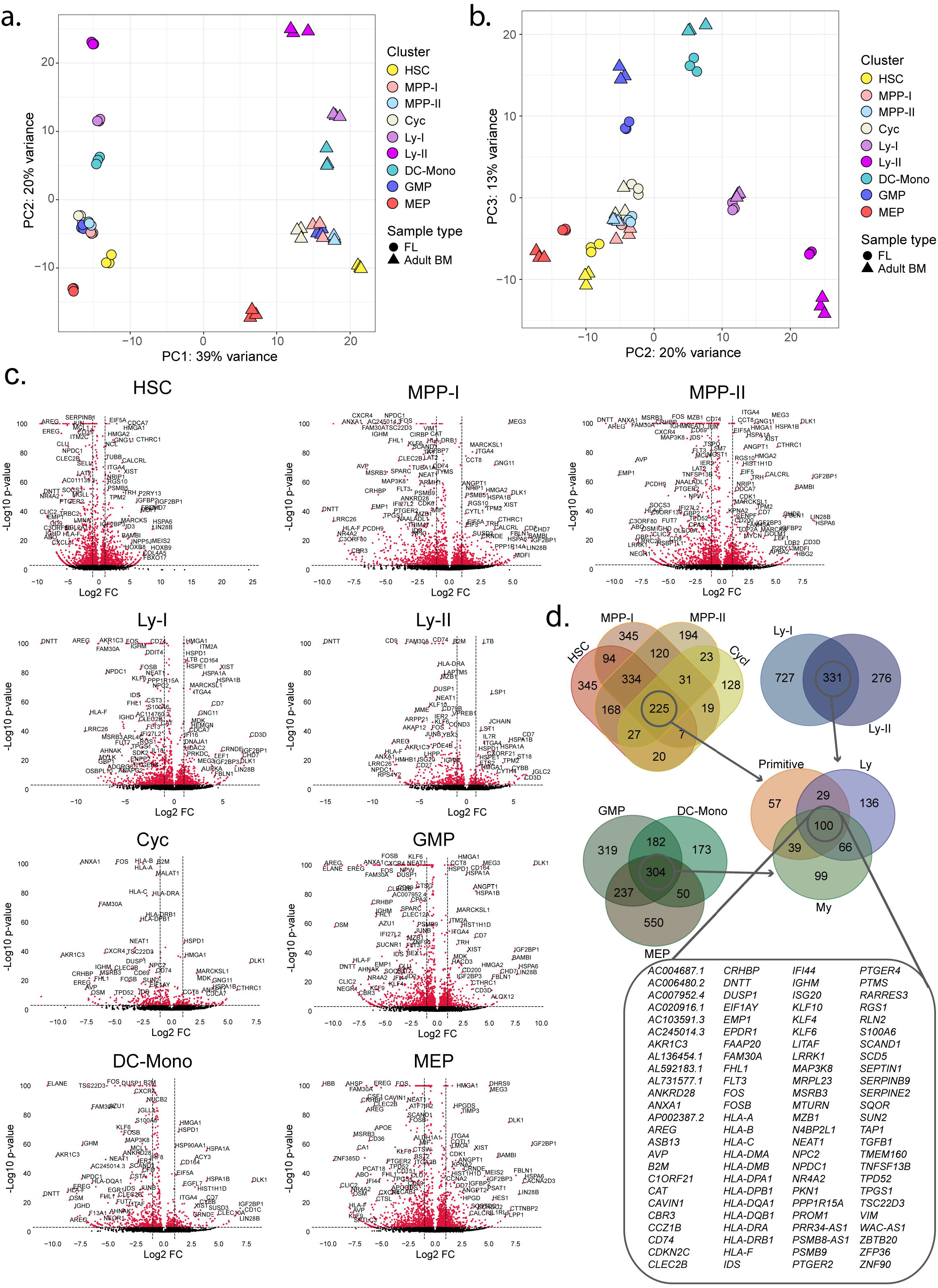
Differential gene expression of CS22 FL and adult BM cells. a-b) PCA of pseudo-bulked clusters from FL CS22 and adult BM, of 500 top variably expressed genes. Colors represent clusters and shapes represents sample type (*circle*: FL CS22 and *triangle*: adult BM). a) PC1 vs PC2; b) PC2 vs PC3. c) Volcano plots of differently expressed genes for pseudo-bulked clusters, x-axis display log2 fold change (FC) and y-axis display −log10 adjusted p-values. Positive FC: enriched in FL, Negative FC: enriched in adult BM. d) Venn diagrams of downregulated genes in FL CS22 compared to adult BM defined by fold change ≤ −2 and an adjusted p-value of <0.05 for each population.

**Sup. Figure 4.**
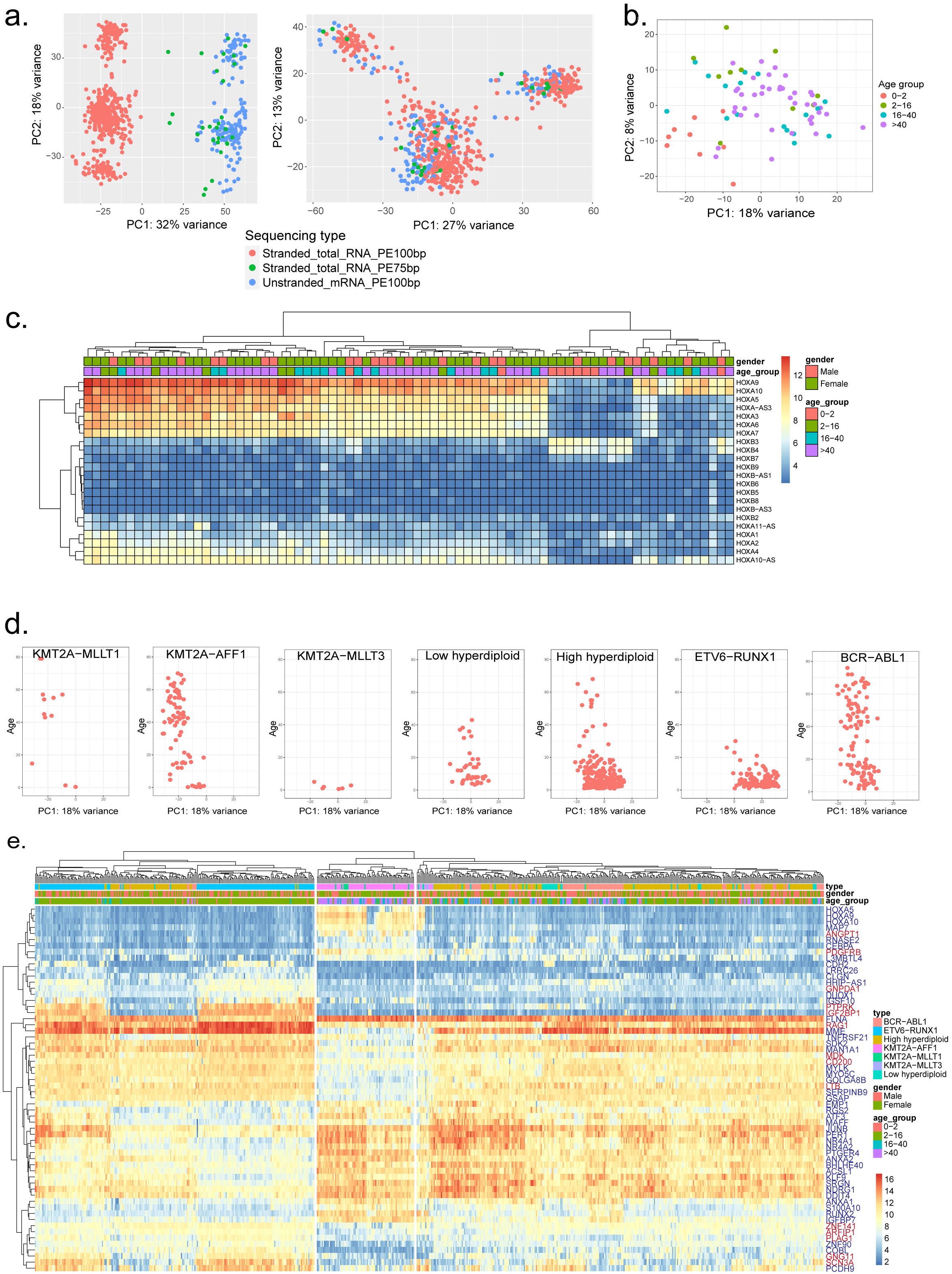
Fetal core signature in B-ALL patient samples. a) B-ALL samples (Gu et al., 2019) before batch correction (*left*) and after batch correction (*right*), showing top 500 variably expressed genes. Color coded according to sequencing method. b) PC1 vs PC2 of MLL-AF4 B-ALL using the fetal core signature. Age group color code is shown to the right. c) Heatmap of *HOX* gene expression in MLL-AF4 ALL. Top rows show gender and age group respectively, color code is shown to the right. d) Each leukemia type from the PCA plot in *Figure 5d* is shown separately. PC1 vs age. e) Heatmap of 60 differentially expressed genes (up or down) separating PC1 for all leukemia types investigated.

## AUTHOR CONTRIBUTIONS

Conceptualization, M.S, C.B and G.K; Methodology, M.S; Formal analysis; M.S.; Data Curation, M.S., P.D. and R.O.; Investigation; M.S., S.P. and R.O.; Visualization, M.S. and C.B; Writing-Original Draft, M.S. and C.B; Writing-Review and Editing, M.S., C.B and G.K.; Funding Acquisition, C.B. and G.K.; Supervision, C.B. and G.K.

## ACKNOWLEDGEMENTS

We would like to acknowledge the personnel of the Lund Stem Cell Center FACS core facility, Anna Fossum and Zhi Ma. Additionally, the Lund University Center for Translational genomics for their assistance with the CITE-seq experiments, especially Linda Geironson Ulfsson and Eva Erlandsson. This work was supported by grants from the Crafoord Foundation, the Swedish Cancer Society, The Ragnar Söderberg Foundation, the Knut and Alice Wallenberg Foundation, the Swedish Research Council, the Swedish Society for Medical Research and the Swedish Childhood Cancer Foundation.

## METHOD

### Sample preparation

Human FLs were donated from elective terminations of pregnancy after informed consent and with the approval of the Ethics Review Authority and the Swedish National Board of Health and Welfare. Fetuses were staged according to Carnegie Staging (CS) and were all from first trimester of pregnancy (developmental stages CS16-9pcw). Single cell suspension of the FL was obtained through mechanical disruption and dissociated through a 40µm filter. The cells were frozen in StemCellBanker (Amsbio) or in Fetal Bovine Serum (FBS) with 10% Dimethyl Sulfoxide (DMSO). Samples were stored at −150°C until day of experiment. CB and adult BM (20-30 years old) data set were from (Sommarin et al., 2021).

### Sample preparation and cell sorting

FL samples were thawed on the day of experiment, washed with Phosphate-Buffered Saline (PBS) with FBS and stained with a panel of CITE-seq antibodies (*Sup. table 2*). In addition each sample was stained with a Hashtag antibody to facilitate sample multiplexing, Fc Receptor blocking reagent (Miltenyibiotec), CD45-Alexa700 (HI30, Biolegend), CD34-FITC (581, Biolegend) and lineage markers (LIN: CD19-BV605 (SJ25C1, BD Bioscience), CD3-PE-Cyanine5 (UCHT1, Biolegend), CD2-PE (RPA-2.10, Biolegend), CD16-BV421 (3G8, Biolegend), CD14-PE-Cyanine7 (M5E2, Biolegend), CD235a-PE-Cyanine5 (GA-R2, BD Bioscience)).These were incubated at 4°C for 30 min, washed and dissolved in PBS+2%FBS (GE lifesciences) with 1/200 7-AAD (BD Bioscience). Up to 20 000 LIN^-^CD45^+^CD34^+^ cells were sorted from each sample using a BD FACSAriaIIu and loaded into the 10x genomics 3’ version 2 platform. The CB and adult BM data sets from (Sommarin et al., 2021) were generated in a similar way from LIN^-^CD45^+^CD34^+^ cells.

### Single-cell CITE-seq library generation and sequencing

After sorting, the samples were loaded on to 10x genomics 3’ version 2 platform (10x Genomics) and single-cell RNA-seq was performed according to the manufacturers instruction with minor changes according to (Stoeckius et al., 2017) to allow for CITE-seq. After revers transcription and cDNA amplification the resulting libraries were sequenced on a NOVAseq (Illumina). Post sequencing the BCL files were processed using cellranger mkfastq to produce FASTQ files, which were then processed using cellranger count to perform alignment to Hg38, filtering, barcode counting and UMI counting. This resulted in the matrix files which were further analyzed in sequential steps using Seurat (Hao et al., 2021) and Scarf (Dhapola et al., 2021).

### Cell filtering and UMAP creation

The out-put from cellranger was loaded into Seurat (Hao et al., 2021) (V4), where the samples were de-multiplexed using the HASH-tag antibody, and cell duplicates were removed. Cells were then filtered based on UMI- and gene counts, to remove low quality cells (*Sup. Table3*). Additionally, UMI counts, cell cycle scores, ribosomal- and mitochondrial contamination were regressed out, using Seurat’s ScaleData function. Each sample was treated separately, RNA reads were LogNormalized and ADT information was CLR (centred-log ratio) normalized using the Seurat NormalizeData function. FL samples from CS22 were integrated into a single data object by Seurat’s IntegrateData function. The output data was then used to make a unified UMAP of the samples. The cells were clustered using the FindClusters function with a resolution of 0.7, which resulted in 11 clusters.

### ADT gating analysis

The CLR normalized ADT count data from Seurat was loaded into Python, were the CLR normalized values were transformed by their antilog (base e) and then multiplied by 1000. The values were then exported to fcs files using the write_FCS function of fcswrite. The FCS files were loaded into FlowJo V10 (BD), where conventional gating was performed, using internal negative controls to set the gates. The gated cells were exported as csv using the export gate option in FlowJo, where upon the gated populations were loaded into Python again. Here the ADT values from cells in the exported gates were matched to the original data to identify the cell identities.

Populations (FL and adult BM) were defined according to the following immunophenotype: HSC^CD49F^: CD34^+^CD38^low^CD45RA^-^CD90^+^CD49F^+^; HSC^CD90^: CD34^+^CD38^low^CD45RA^-^ CD90^+^; MPP: CD34^+^CD38^low^CD45RA^-^CD90^-^; LMPP: CD34^+^CD38^low^ CD45RA^+^CD90^-^; CMP^CD123^: CD34^+^CD38^+^CD10^−^CD45RA^-^CD123^+^, CMP^CD135^: CD34^+^CD38^+^CD10^−^CD45RA^-^ CD135^+^; GMP ^CD123^: CD34^+^CD38^+^CD10^−^CD45RA^+^CD123^+^; GMP^CD135^ CD34^+^CD38^+^CD10^−^CD45RA^+^CD135^+^; MEP ^CD123^: CD34^+^CD38^+^CD10^−^CD45RA^-^CD123^-^; MEP^CD135^: CD34^+^CD38^+^CD10^−^CD45RA^-^CD135^-^; CLP^CD10^: CD34^+^CD38^+^CD45RA^+^CD10^+^, CLP^IL7R^: CD34^+^CD38^+^CD45RA^+^IL7R^+^; CLP^CD7^: CD34^+^CD38^+^CD45RA^+^CD7^+^.

### Projection of samples onto reference UMAP

Projection of cells onto the FL CS22 reference UMAP was done by using Scarf (Dhapola et al., 2021). In brief the UMAP coordinates, HVGs and cluster information of the FL CS22 analysis in Seurat was loaded into Scraf, and by using the run_mapping function (with k=11) the nearest neighbors of each cell projected was calculated. This was performed for FL CS16, FL 9 pcw, CB and adult BM, the latter two from (Sommarin et al., 2021). The same analysis was performed using the adult BM as a reference, with the same settings as for the FL CS22 and then projecting FL CS16, FL CS22, FL 9pcw and CB cells. To classify each projected cell to a cluster the get_target_classes function (threshold = 0.4) was used. The mapping score was calculated using get_mapping_score, and the size of each cell in the reference map was set proportional to the mapping score. The same analysis was performed using data from (Roy et al., 2021).

### Investigation of molecular differences in primitive FL sub-clusters

To define the molecular differences between the HSC, MPP-I and MPP-II within FL CS22 the FindMarkers function of Seurat was used for each individual cluster (adjusted P value <0.001, log2 fold change (FC) >|0.5|). These marker genes together with the cells from the HSC, MPP-I and MPP-II clusters were then used to make a subsetted cell-gene-matrix of the scarf normalized values. Next, hierarchical clustering was performed using the clustermap function from the seaborn package. The resulting dendrogram of genes were cut into nine clusters.

### CellRadar

To define the lineage affiliation of each gene cluster, the BloodSpot dataset ‘normal human hematopoiesis’ was used (Bagger et al., 2016). A radar plot was generated using min-max scaled median value of marker genes in each cluster. CellRadar (*Dhapola et al. manuscript in preparation*, available here: https://github.com/KarlssonG/cellradar)

### Differential gene expression testing

To perform differential gene expression analysis, each cluster of the FL CS22 and the adult BM cells predicted to a FL cluster were pseudo-bulked into three replicates for each sample and cluster, using the make_bulk function of Scarf (Dhapola et al., 2021), this was additionally performed on the ADT gated immunophenotypic populations. These pseudo-bulk populations were then exported as csv and loaded into DEseq2 (Love et al., 2014). DEseq2 then performed differential gene expression comparing each cluster of FL CS22 to its corresponding predicted cluster in adult BM. Here, clusters DC-I and Ly-III were excluded in the analysis due to their low cell count (<60 cells). The following result file were then analyzed for up-(FL signal) and down-(adult BM signal) regulated genes using the adjusted p-value <0.05 and FC ≥|2| to make gene lists for each cluster, visualized in volcanoplots by matplotlib.pyplot. Additionally, the DEseq2 normalized values were exported for each cluster and loaded into GSEA (Subramanian et al., 2005) to analyze hallmark gene sets (Liberzon et al., 2015). Gene sets with an FDR q-value of <0.05 were considered significantly enriched/depleted (nr of permutations:1000, permutation type: gene-set).

### Principal component analysis

To perform PCA on the clusters and the ADT-defined immunophenotypic populations the pseudo-bulked data was loaded into DEseq2. Again, the DC-I and Ly-III clusters were excluded due to their low cell count. Here, the data was transformed using the VST function of DEseq2, after which the top 500 most variable genes were used in the prcomp function of R to perform PCA. Finally, the data was visualized using ggplots2.

### Definition of fetal derived gene signature

The significant genes (FC ≥|2| and adjusted p-value of <0.05) for each cluster of the DEseq2 differential testing were analyzed using the Venn tool of (http://bioinformatics.psb.ugent.be/webtools/Venn/). The clusters HSC, MPP-I, MPP-II and Cyc were first compared to find which genes were in common. Next the clusters MEP, GMP and DC-Mono were compared, and finally the Ly-I and Ly-II clusters. Clusters DC-I and Ly-III were excluded in differential gene expression analysis due to their low cell count (<60 cells). Finally, the genes in common between the three comparisons were linked to find a strict list of genes in common for all clusters. The gene names of these were exported as csv, for further analysis.

### Analysis of fetal derived gene signature in FL and adult BM

To investigate the molecular differences between FL and adult BM within the single cell data, the BM and FL datasets were combined by using the ZarrMerge function of Scarf. To analyze the differences in HOX usage in the clusters, all HOX genes expressed in >10 cells were used to create a heatmap using the plot_marker_heatmap function of scarf. To investigate the expression of the fetal derived gene signature, the genes specific for FL was used to subset the combined FL-BM dataset. Next, using the clustermap function of seaborn the cells of the FL-BM dataset were clustered using hierarchical clustering (Euclidean distances and Ward linkage).

### Investigation of the fetal derived gene signature in pediatric ALL

To investigate the fetal gene signature in the ETV6-RUNX1 IPSC model publicly available data from (Boiers et al., 2018) were downloaded. FPKM values were log-transformed using log2 (FPKM +1) and then scaled and centered using the scale function.

To investigate the fetal gene signature in childhood ALL, a publicly available data set of approximately 2000 patients was used (Gu et al., 2019). From this data set we focused on patients with the following cytogenetic abnormality: KMT2A-AFF1, KMT2A-MLLT1, KMT2A-MLLT3, BCR-ABL1, ETV6-RUNX1 and high or low hyperdiploidy. The HTSeq files of these were loaded into DEseq2 using the DESeqDataSetFromHTSeqCount function. Next the counts function were used to get a DEseq2 dataframe. The counts were transformed using variance stabilizing transformation by the VST function, and batch effects removed with LIMMA::removeBatchEffect. Thereafter the gene lists from DE-analysis of FL and adult BM were loaded in and using the getBM function of Biomart was used to translate the gene names into hg37 ensembl IDs. The FL and adult BM specific ensembl IDs were used to subset the transformed data, selecting only our genes of interest, excluding gender associated genes. PCA was performed on the KMT2A-AFF1 translocation using the prcomp function, calculating all principal components. PC1 was shown to capture the age differences in the samples, and thus the top 25 genes and the lowest 25 genes from PC1 were used generate a heatmap of the samples. To analyze all samples of interest PCA was again used, following the same procedure as previously described. This resulted in another set of PCs, where again PC1 described the age-related changes. Here the top 30 and the lowest 30 genes were used to make the heatmap.

**Sup. Table 1.**
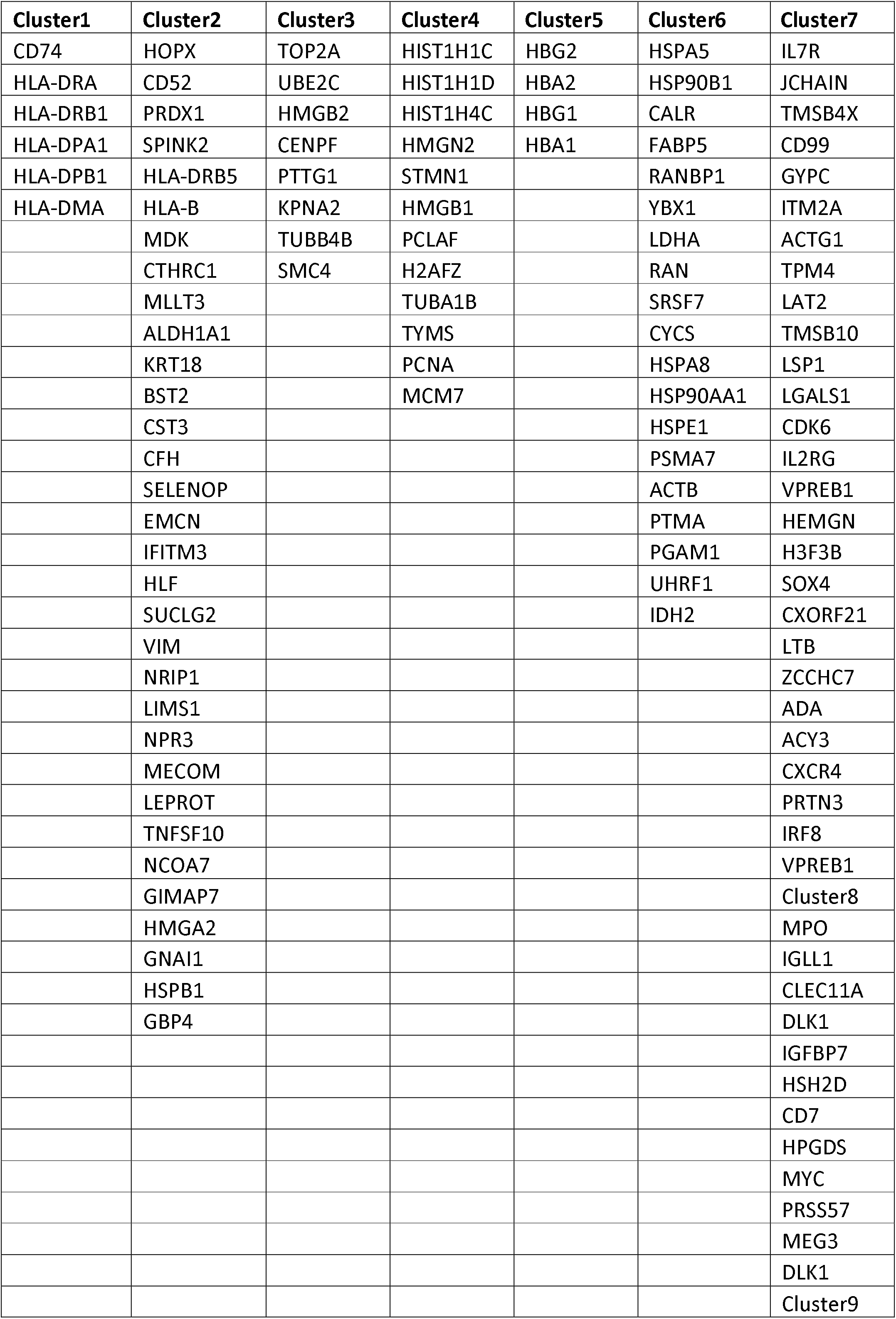
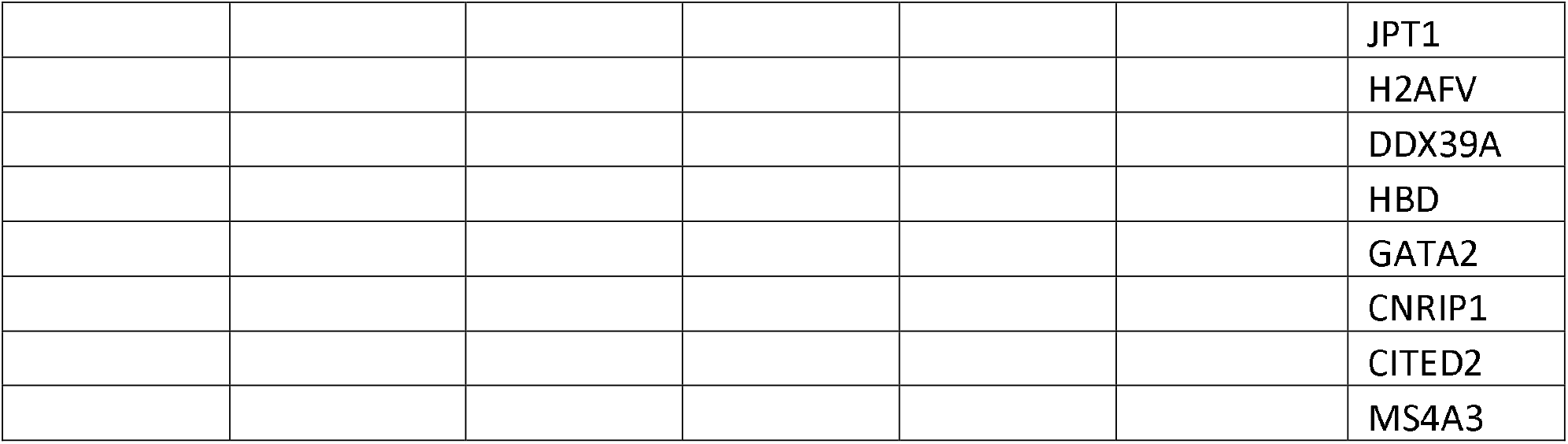
Gene list shown per cluster from heatmap in Figure 3e.

**Sup. Table 2:**
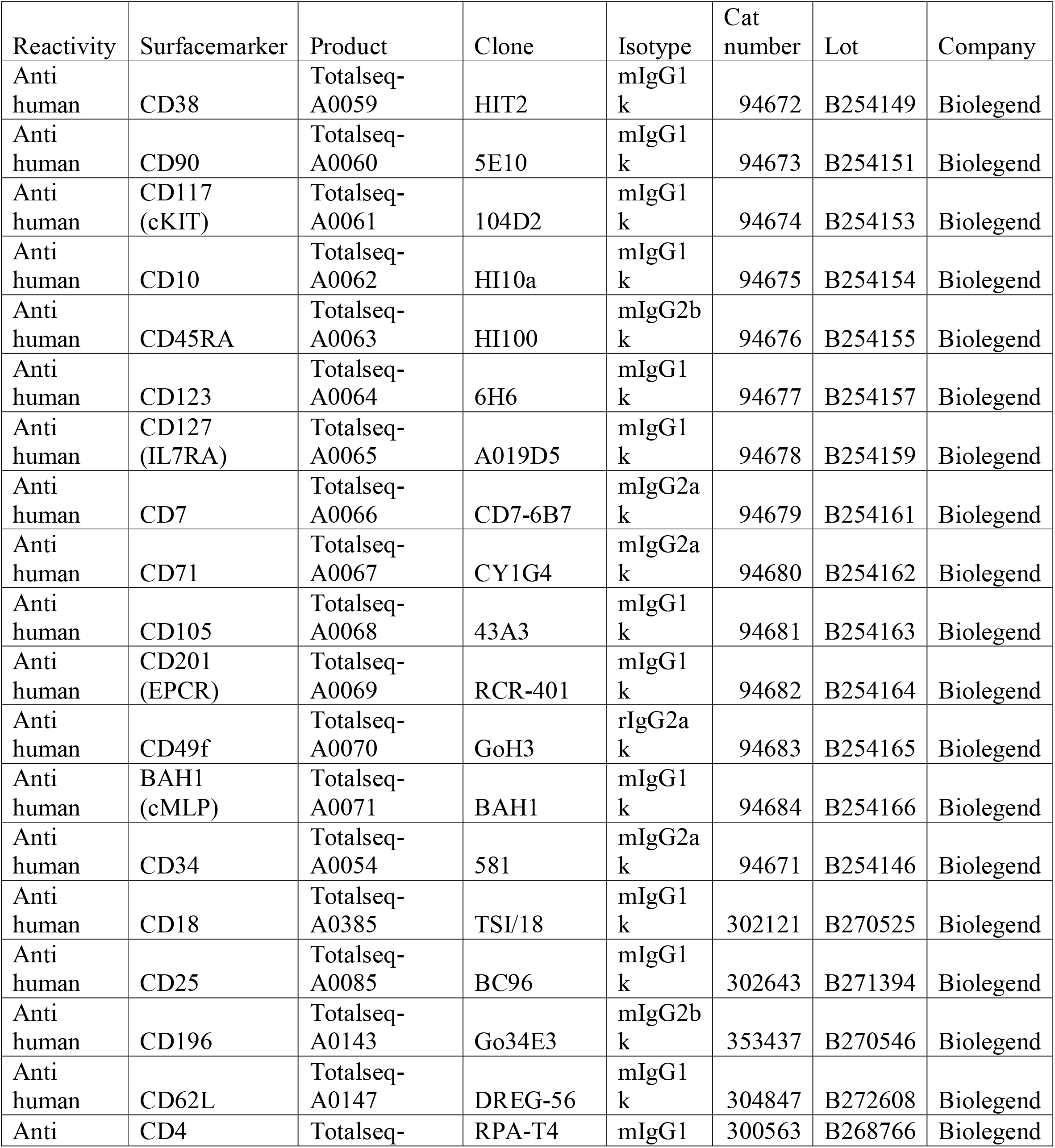
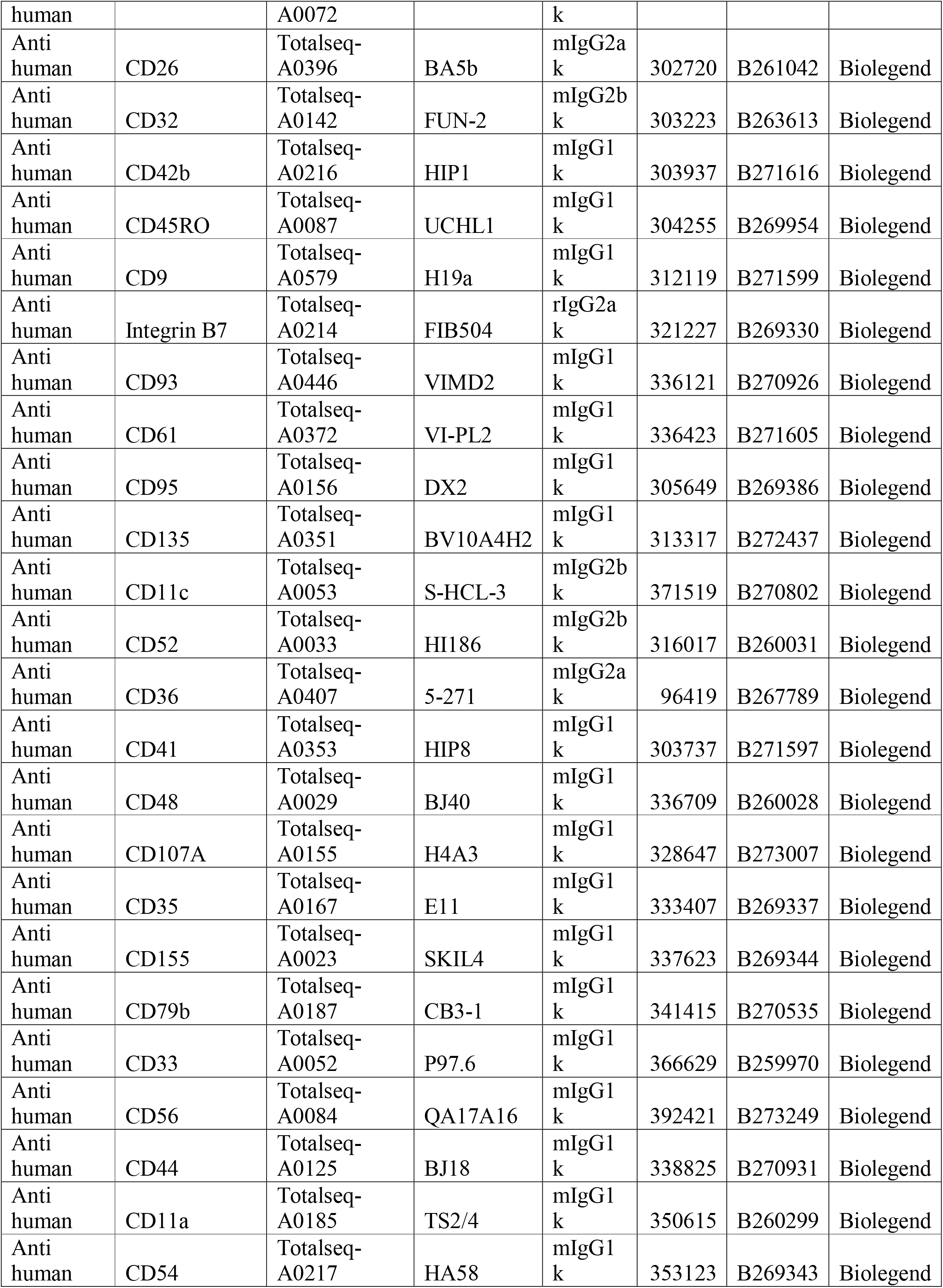
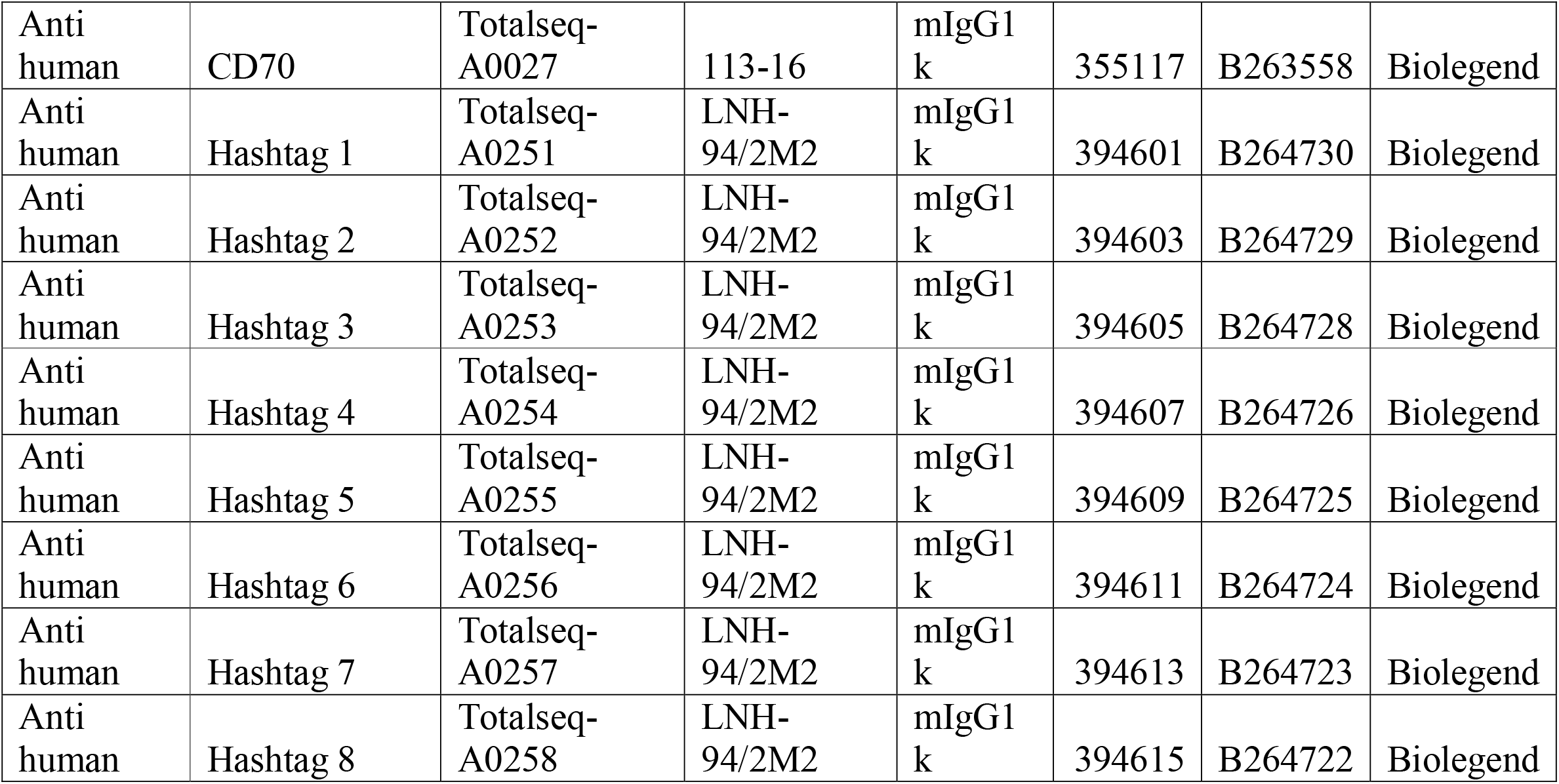
CITE-seq antibodies used.

**Sup. Table 3:** Samples included in the study.

| Sample | # Sorted cells | # Cells used in analysis |
| --- | --- | --- |
| FL CS22 sample1 | 10 000 | 3040 |
| FL CS22 sample2 | 10 000 | 3692 |
| FL CS16 sample1 | 906 | 109 |
| FL CS16 sample2 | 244 | 34 |
| FL 9pcw sample1 | 2 213 | 297 |
| FL 9pcw sample2 | 4 946 | 842 |

